# Genomic DNA preparation enabling multiple replicate reads for accurate nanopore sequencing

**DOI:** 10.1101/035436

**Authors:** Dimitra Tsavachidou

## Abstract

Sequencing at single-nucleotide resolution using nanopore devices is performed with reported error rates 10.5–20.7% (Ip et al., 2015). Since errors occur randomly during sequencing, repeating the sequencing procedure for the same DNA strands several times can generate sequencing results based on consensus derived from replicate readings, thus reducing overall error rates.

The method presented in this manuscript constructs copies of a nucleic acid molecule that are consecutively connected to the nucleic acid molecule. Such copies are useful because they can be sequenced by a nanopore device, enabling replicate reads, thus improving overall sequencing accuracy.

## Serial Copies Method

Long-read sequencing technologies can be benefited significantly by sample preparation methods that enable multiple replicate reads of the same genomic DNA molecule. For example, the PacBio (Pacific Biosciences) sequencing error rate for a single read is relatively high (around 11%-15%) (Rhoads and Au, 2015). The Circular Consensus Sequencing (CCS) method allows for the repeated sequencing of individual templates (Travers et al., 2010). The errors are distributed randomly in single reads, so that the overall error rate can be reduced by generating CCS reads with sufficient sequencing passes. For example, templates with at least 4 replicate reads (i.e. templates that went through at least 4 CCS passes) have a minimum Phred quality score of 20, templates with 7 replicate reads have a Phred score of at least 40, and those with 9 replicate reads have a Phred score of at least 50 (Larsen et al., 2014).

The MinION nanopore sequencing platform (Oxford Nanopore Technologies) can generate 2D reads (one read for the template and one for its complement), reducing the error rate to approximately 12% (Ip et al., 2015).

Clearly, there is a need for further error rate reduction in nanopore sequencing. Methods of sample preparation that generate multiple DNA copies in tandem are desired. One such method (called the “serial copies method”) is shown in Figure 1.

During the 1st step of the method, a blunt-ended doubled-stranded DNA molecule is ligated to two adaptors, one of which is a hairpin. The other adaptor can be free in solution or attached to a solid surface. During the 2nd step, the DNA molecule is subjected to incubation with nicking endonucleases that recognize a restriction site within the adaptor. The nicking endonucleases may nick within the adaptor or between the 3’ end of the adaptor and the adjacent 5’ end of the DNA molecule.

**Figure 1:**
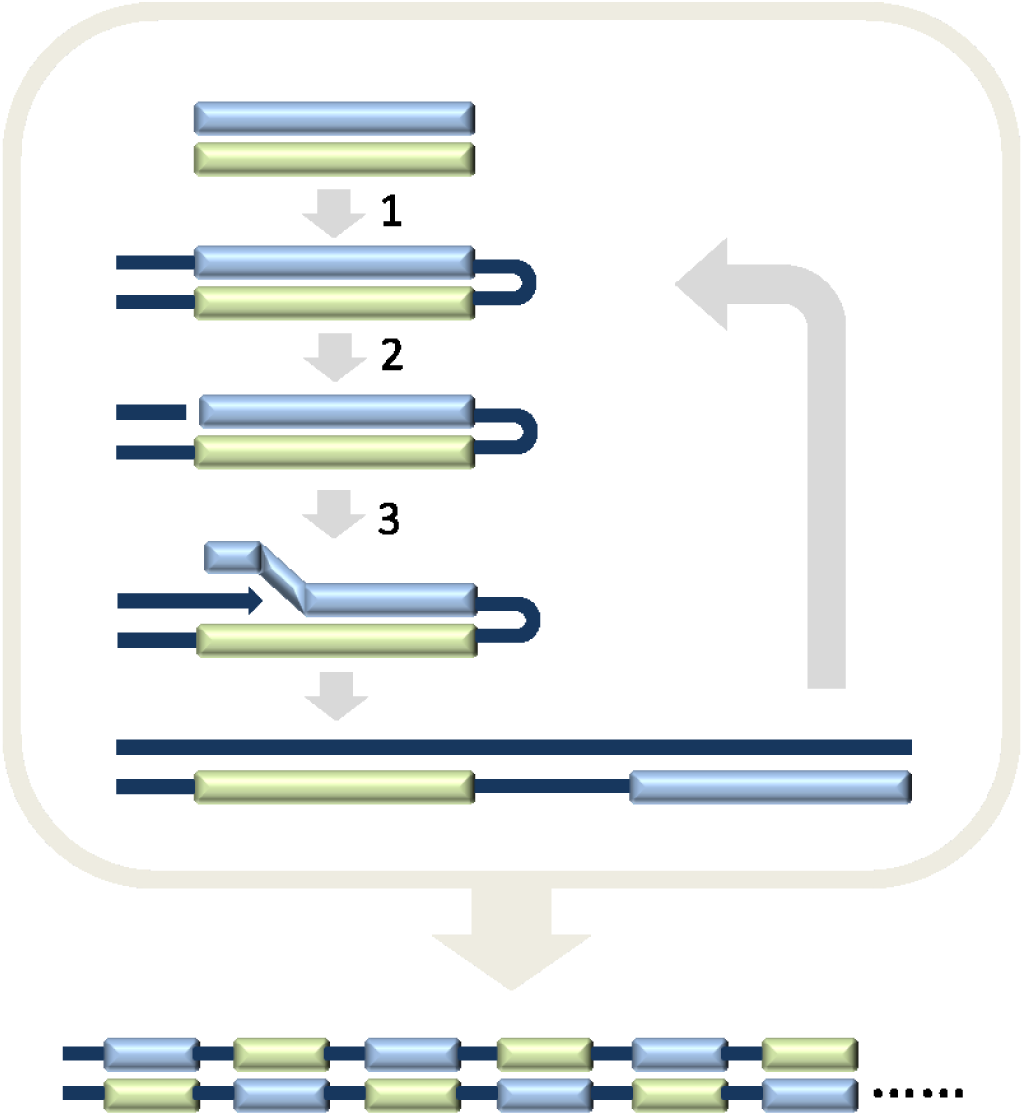
Serial copies method

During the 3rd step, the DNA molecule participates in an extension reaction using strand-displacing polymerases. Owing to the presence of the hairpin, two connected copies of the genomic DNA are generated, one inverted in relation to the other.

The process can be repeated several times by subjecting the product of each extension reaction to a cycle of hairpin adaptor ligation, nicking and polymerization. The total number of copies generated after each cycle is double the number of copies of the previous cycle.

The design of this method allows running all steps concurrently, by including ligases, hairpin adaptors, nicking endonucleases and strand-displacing polymerases in a single reaction.

Importantly, the single reaction is combined with a proprietary procedure running concurrently, that prevents the formation of hairpin-to-hairpin ligation products and removes any such products formed, without using laborious and expensive size selection procedures (e.g. AMPure, gel separation). Additional information and data about this approach will be presented in future revisions of this manuscript.

The size of the construct produced by the serial copies method depends on the type of polymerase used in the extension steps. When phi29 is used, the total size can be ~50 kb. The smaller the original genomic fragment to be copied, the more copies can be generated for a fixed total construct size.

Nicking endonuclease sites may be present in the genomic fragments to be copied using the serial copies method. In this event, genomic fragments may be nicked and bias may be introduced, because of underrepresentation of regions at the 5’ sides of nicks. Such bias may be prevented by performing two separate reactions for the same genomic sample, using a different type of nicking endonuclease in each reaction. There are other versions of the serial copies method that do not require nicking endonucleases. These versions will be discussed in future revisions of this manuscript.

An advantage of this method is that it produces multiple copies of a genomic fragment in a convenient double-stranded construct that can be subsequently attached to appropriate adaptors for nanopore sequencing. The serial copies method can be more useful than rolling-circle amplification, which produces less convenient single-stranded constructs and requires circularization which is typically an inefficient process. For example, circularization is used in the construction of mate-pair libraries, an expensive process that may require up to 15–20 μg of high molecular weight DNA of which most is lost during the enrichment step of the end-to-end ligated fragments (Knief, 2014). The widely used Nextera mate pair library preparation kit (Illumina) requires a minimum of 1 ug starting material to yield 2–8 kb products, but getting longer fragments (5–10 kb) requires more material (4 ug) and sacrifices library diversity (Illumina, 2014).

